# Bacterial Membrane Vesicles in Wastewater Disseminate Antibiotic Resistance Genes

**DOI:** 10.64898/2026.05.11.724380

**Authors:** Hongyue Zhang, Mahmud Syed, Zhenzhen He, Xiaonan Tang, Rui Fu, Yujie Men, Tiong Gim Aw, Joan B. Rose, Danmeng Shuai, Yun Shen

## Abstract

Bacterial membrane vesicles (BMVs) have emerged as important contributors to the dissemination of antibiotic resistance genes (ARGs) in the environment. Here, we developed a high-performance immunomagnetic isolation method that improves the purity and selectivity of BMV recovery from wastewater, minimizes contamination from eDNA and viruses, and enables differentiation of BMVs originating from Gram-positive versus Gram-negative bacteria with minimal cross-reactivity. Using this approach, we found that ARGs such as the kanamycin resistance gene (*kan*^*R*^) was highly abundant in BMVs from both raw and treated wastewater, exhibited persistence following treatment, and retained the ability to generate antibiotic-resistant bacteria via transformation. Metagenomic sequencing further revealed that tetracycline resistance genes were the most abundant ARG class across all wastewater samples, while the composition of BMV-associated ARGs differed from the bulk ARG profile. These findings highlight the critical yet underrecognized role of BMVs in the spread of antimicrobial resistance and underscore the need to address BMV-mediated pathways within a One Health framework linking environmental and human health.

## Main

Spread of antimicrobial resistance in bacteria increases the mortality of infection, posing a global public health challenge. In 2021, bacterial antimicrobial resistance directly resulted in 1.14 million deaths worldwide and were associated with 4.71 million deaths^1^. The environmental dissemination of antibiotic resistance genes (ARGs) is an important driver of antimicrobial resistance. Especially, wastewater is an ARG reservoir and often contains ARG concentrations orders of magnitude higher than natural environments^2^. The high density and diversity of bacterial communities in wastewater treatment processes can facilitate horizontal gene transfer (HGT), which enables the transfer of ARGs among bacteria, drives pathogen evolution, and accelerates the spread of antimicrobial resistance^3–6^. Many treatment processes inadequately remove ARGs^4,7–10^, and aerobic biological treatment can even increase their abundance^3,10,11^. As a result, ARGs in treated effluents and biosolids can spread through water reuse and land application, threatening food safety, ecosystems, and human health^12^. Therefore, elucidating the fate and dissemination of ARGs in wastewater treatment is critical for informing effective antimicrobial resistance control strategies.

While conjugation, transformation, and transduction were long recognized as the primary HGT mechanisms, the transfer of ARGs through bacterial membrane vesicles (BMVs) was recently discovered as an emerging pathway^13–16^. BMVs are 10-500 nm lipid-bilayer particles released by Gram-positive and Gram-negative bacteria^14^ that carry nucleic acids, proteins, and other biomolecules, mediating HGT, intercellular communication, cargo delivery, and adaptation to environmental stress^17^. BMV-mediated transfer of ARGs from *Escherichia coli* O157:H7 to *Salmonella enterica* was found to be more efficient than natural transformation^16^. In addition, ARGs protected inside a membrane structure may exbibit enhanced environmental stability and resistance to stressors such as heat treatment and enzymatic digestion^18,19^. It is increasingly recognized that BMVs may play a significant yet underexplored role in the spread of antimicrobial resistance; however, our understanding of the environmental behavior of BMV-associated ARGs is limited, hampering the development of effective control strategies.

Although BMV-associated ARGs have been detected in wastewater^20–22^, their fate and removal across wastewater treatment processes remain poorly characterized. One critical knowledge gap is the lack of methods for selectively isolating BMVs from complex matrices without co-isolating impurities such as extracellular DNA (eDNA) and free bacteriophages, which can also carry ARGs. Differential and density-gradient ultracentrifugation are the most commonly used methods for BMV isolation, often complemented by filtration and size-exclusion approaches^23–27^. However, these size- and density-based techniques can lead to significant co-isolation of contaminants, including bacteriophages^25^. In addition, vesicles can protect their internal cargo (e.g., biomolecules and viruses) from disinfection and other environmental stressors^28–31^. However, little is known about the response and removal of BMV-associated ARGs during wastewater treatment. In particular, the extent to which BMV-associated ARGs contribute to overall ARG persistence and dissemination during wastewater treatment, and their capacity to mediate HGT, remains largely unexplored.

To address these knowledge gaps, we developed and applied a selective immunomagnetic isolation approach to investigate BMV-associated ARGs across a full-scale wastewater treatment system. Specifically, we leveraged surface biomarkers unique to Gram-positive and Gram-negative bacteria to enable high-purity and origin-specific recovery of BMVs from complex wastewater matrices while minimizing interference from eDNA and viruses. Using this approach, we quantified BMV-associated ARGs along the treatment train and evaluated their persistence relative to the total ARG pool. Furthermore, we evaluate BMV-mediated HGT using transformation assays and characterize ARG composition through metagenomic analysis. Our results reveal that BMVs serve as persistent, selective, and functional carriers of ARGs, exhibiting resistance to conventional treatment processes and becoming increasingly dominant in treated effluents. This work advances our understanding of antibiotic resistance dynamics in wastewater systems, highlights the need to incorporate vesicle-mediated processes into risk assessment, and supports the design of next-generation treatment strategies for safe water reuse.

### Immunomagnetic isolation enables selective and high-purity recovery of bacterial membrane vesicles from wastewater

BMVs released from both Gram-positive and Gram-negative bacteria^32,33^ were harvested from antibiotic-resistant laboratory cultures (i.e., methicillin-resistant *Staphylococcus aureus* (MRSA) and *Escherichia coli*) for method development, and from municipal wastewater to validate the method in environmental samples (**Figs. 1a and S1**). Our unique contribution is the application of immunomagnetic isolation using lipoteichoic acid (LTA)^34^ and lipopolysaccharide (LPS)^35^ antibodies to selectively capture Gram-positive and Gram-negative BMVs, respectively, as LTA and LPS are well-established surface biomarkers of Gram-positive and Gram-negative bacterial cell walls and their BMVs^36^. We first evaluate the capacity of immunomagnetic BMV isolation. *E. coli*-derived BMVs were purified by immunomagnetic selection, then spiked into buffer and re-isolated using the same method, yielding near-complete recovery (94.45%). MRSA- and *E. coli*-derived BMVs were also obtained via ultracentrifugation, spiked into particle-depleted raw wastewater, and recovered by immunomagnetic isolation, with efficiencies of 51.89% for Gram-positive BMVs and 45.65% for Gram-negative BMVs (**Fig. 1b**). These efficiencies were substantially higher than those achieved by ultracentrifugation alone, which recovered only 8.53% and 5.67% of spiked BMVs from particle-depleted wastewater and phosphate-buffered saline (PBS), respectively (**Fig. S2**). These results demonstrate that immunomagnetic isolation enables efficient BMV recovery in complex wastewater matrices, with efficiencies suitable for quantitative analysis and exceeding those of ultracentrifugation and previously reported yields in complex human body fluids^37,38^.

**Fig. 1.**
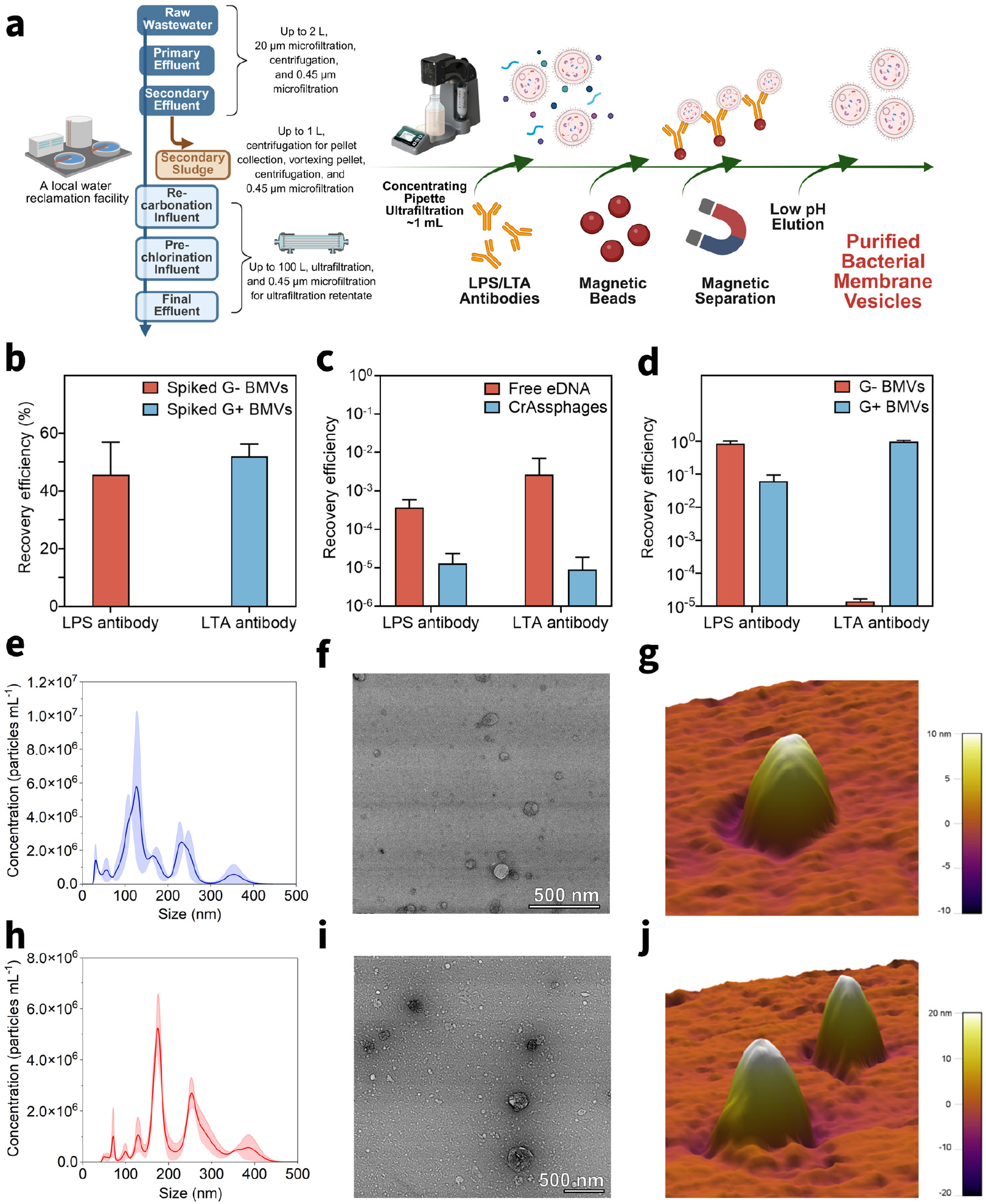
Immunomagnetic isolation of bacterial membrane vesicles (BMVs) from laboratory cultures and wastewater. **a**, Schematic diagram of sample collection and vesicle isolation from a water reclamation facility. Lipoteichoic acid (LTA) and lipopolysaccharide (LPS) antibodies were used for Gram-positive (G+) and Gram-negative (G-) BMV isolation, respectively. **b**, Recovery efficiency of BMVs by immunomagnetic isolation method. BMVs derived from laboratory-cultured methicillin-resistant *Staphylococcus aureus* (MRSA) harboring plasmid pUSA300-HOU-MR and *E. coli* harboring plasmid pCM184 were spiked into particle-depleted raw wastewater and subsequently isolated. **c**, Recovery efficiency of plasmid pCM184 and crAssphages by immunomagnetic isolation method. pCM184 was spiked into particle-depleted raw wastewater and then isolated, while endogenous crAssphage was directly recovered from untreated raw wastewater. **d**, Cross-reactivity of immunomagnetic isolation method. LTA or LPS antibodies were separately incubated with phosphate-buffered saline containing BMVs derived from MRSA (G+) or *E. coli* (G-). Recovery efficiency in **b-d** was determined as the ratio of isolated gene copy concentrations to their initial concentrations, as quantified by quantitative polymerase chain reaction. Data are presented as mean ± SD (n=3-4). **e**, Nanoparticle tracking analysis (NTA) of raw wastewater-derived G+ BMVs (n=3). **f**, Transmission electron microscopy (TEM) of raw wastewater-derived G+ BMVs. Scale bar = 500 nm. **g**, Atomic force microscopy (AFM) topography of raw wastewater-derived G+ BMVs. **h**, NTA of raw wastewater-derived G-BMVs (n=3). **i**, TEM of raw wastewater-derived G-BMVs. Scale bar = 500 nm. **j**, AFM topography of raw wastewater-derived G-BMVs.

We next evaluated the selectivity of immunomagnetic BMV isolation. Free plasmids representing eDNA (pUSA300HOUMR and pCM184 from MRSA and *E. coli*, respectively) were spiked into particle-depleted raw wastewater and subjected to immunomagnetic isolation. Only 0.269% and 0.0372% of plasmids were recovered via LTA- and LPS-based isolation, respectively (**Fig. 1c**). Endogenous crAssphage in untreated raw wastewater was also selected as a representative and abundant bacteriophage to evaluate the selectivity of BMV isolation. Only 9.00 × 10^−4^% and 1.31 × 10^−3^% of crAssphage were recovered via LTA- and LPS-based isolation, respectively (**Fig. 1c**). This result contrasts sharply with sequential differential and density-gradient ultracentrifugation, which co-isolates ~15% of viruses with vesicles from environmental samples^25^. In addition, we evaluated the specificity of immunomagnetic BMV isolation according to their bacterial origin. MRSA- and *E. coli*-derived BMVs obtained via ultracentrifugation were spiked into PBS and recovered using LTA- and LPS-based isolation, respectively, achieving 98.09% and 87.01% recovery for Gram-positive and Gram-negative BMVs (**Fig. 1d**). In contrast, cross-reactivity was minimal, with only 6.08% of Gram-negative BMVs captured by LTA antibodies and 0.00142% of Gram-positive BMVs co-isolated by LPS antibodies (**Fig. 1d**). Together, these results demonstrate that immunomagnetic isolation effectively minimizes interference from eDNA and viruses, enables discrimination of BMVs based on their bacterial origin, and establishes a robust foundation for selective BMV recovery from complex wastewater matrices.

Particle size, morphology, and mechanical properties of purified BMVs from raw wastewater were characterized using nanoparticle tracking analysis (NTA), transmission electron microscopy (TEM), and atomic force microscopy (AFM). Immunomagnetic isolation yielded dominant particle sizes of 127 and 232 nm for Gram-positive BMVs and 175 and 253 nm for Gram-negative BMVs (**Figs. 1e and 1h**). In contrast, concentrated raw wastewater exhibited multiple dominant particle size populations (**Fig. S3**). TEM confirmed the structural integrity and high purity of the isolated BMVs, with no detectable bacterial cells or bacteriophages (**Figs. 1f and 1i**). AFM provided complementary morphological characterization in the liquid phase (**Figs. 1g and 1j**)^39^, revealing mean heights of 30 and 34 nm for wastewater-derived Gram-positive and Gram-negative BMVs, respectively (*p* > 0.05, **Fig. S4a**), and mean lateral sizes of 119 and 189 nm (*p* < 0.001, **Fig. S4b**). The smaller height relative to lateral size likely reflects vesicle flattening due to strong electrostatic interactions with the poly-L-lysine (PLL)-coated mica substrate during AFM characterization^40,41^. AFM further quantified the mechanical properties, showing Young’s modulus values of 671 and 348 MPa for Gram-positive and Gram-negative BMVs, respectively (*p* < 0.01, **Fig. S5**). Together, NTA and AFM results indicate that Gram-negative BMVs are larger, whereas Gram-positive BMVs exhibit greater stiffness. These observations are consistent with previous findings: in Gram-positive bacteria, the cytoplasmic membrane must protrude through a thick peptidoglycan layer, which may favor the formation of smaller BMVs, whereas in Gram-negative bacteria, the outer membrane may more readily bleb off to produce larger BMVs^17^. Gram-positive BMVs are also structurally more rigid, likely due to the presence of cross-linking components (e.g., LTA) and residual peptidoglycan, while Gram-negative BMVs consist primarily of a thinner phospholipid bilayer and are therefore more flexible^42–44^. We further compared the height, lateral size, and stiffness of MRSA- and *E. coli*-derived BMVs with Gram-positive and Gram-negative BMVs isolated from wastewater (**Figs. S4 and S5**). The results highlight the diversity of BMVs in environmental samples, which exhibit distinct size distributions and mechanical properties compared to laboratory-derived BMVs. Nevertheless, the overall trends remain consistent, with Gram-negative BMVs being larger and Gram-positive BMVs exhibiting greater stiffness. The clear distinction in both size and mechanical properties further support the validity of immunomagnetic isolation in selectively capturing two physiologically distinct BMV populations.

### ARGs associated with bacterial membrane vesicles persist throughout wastewater treatment

Kanamycin resistance (*kan*^*R*^), tetracycline resistance (*tet*^*R*^), ampicillin resistance (*amp*^*R*^), and β-lactamase (*bla*^*Z*^) genes were all identified in raw wastewater by quantitative polymerase chain reaction (qPCR); *kan*^*R*^ was abundant and consistently detected and was used as the reference (**Fig. S6**). During conventional wastewater treatment (**Fig. 1a**), total *kan*^*R*^ decreased from 7.84 log_10_ gene copies per liter (log_10_ gc L^−1^) in raw wastewater to 6.79 log_10_ gc L^−1^ in secondary effluent (*p* > 0.05, **Fig. 2a**). In contrast, *kan*^*R*^ associated with Gram-positive and Gram-negative BMVs remained stable (3.98-4.33 log_10_ gc L^−1^ and 4.49-4.27 gc L^−1^, respectively; both *p* > 0.05). Upon transition to advanced treatment, total *kan*^*R*^ dropped sharply to 4.42 log_10_ gc L^−1^ after quick lime flocculation and clarification (*p* < 0.0001 for re-carbonation influent versus secondary effluent) and further declined to 4.03 log_10_ gc L^−1^ in the final effluent. Gram-positive and Gram-negative BMV-associated *kan*^*R*^ showed a similar decrease after chemical flocculation (2.20 and 2.44 log_10_ gc L^−1^, respectively; both *p* < 0.0001 for re-carbonation influent versus secondary effluent) and then stabilized at 2.18 and 2.33 log_10_ gc L^−1^ in the final effluent, respectively. Primary and secondary treatment achieved minimal removal of BMV-associated *kan*^*R*^ (<0.30 log_10_), compared with ~1 log_10_ reduction in total *kan*^*R*^, indicating limited effectiveness against membrane-protected ARGs. When Gram-positive and Gram-negative BMVs were evaluated collectively, BMV-associated *kan*^*R*^ decreased by 2.04 log_10_ across the full treatment train, which was 1.77 log_10_ less than total *kan*^*R*^, also highlighting vesicle-mediated protection and persistence. Quick lime flocculation substantially reduced both total and BMV-associated *kan*^*R*^, likely due to elevated pH (~11) destabilizing vesicle membranes, damaging DNA, and promoting flocculation and precipitation of ARGs^45^. In addition, removal efficiencies from raw wastewater to final effluent were similar for Gram-positive and Gram-negative BMVs (2.00 versus 2.31 log_10_), suggesting persistence is independent of bacterial origin.

**Fig. 2.**
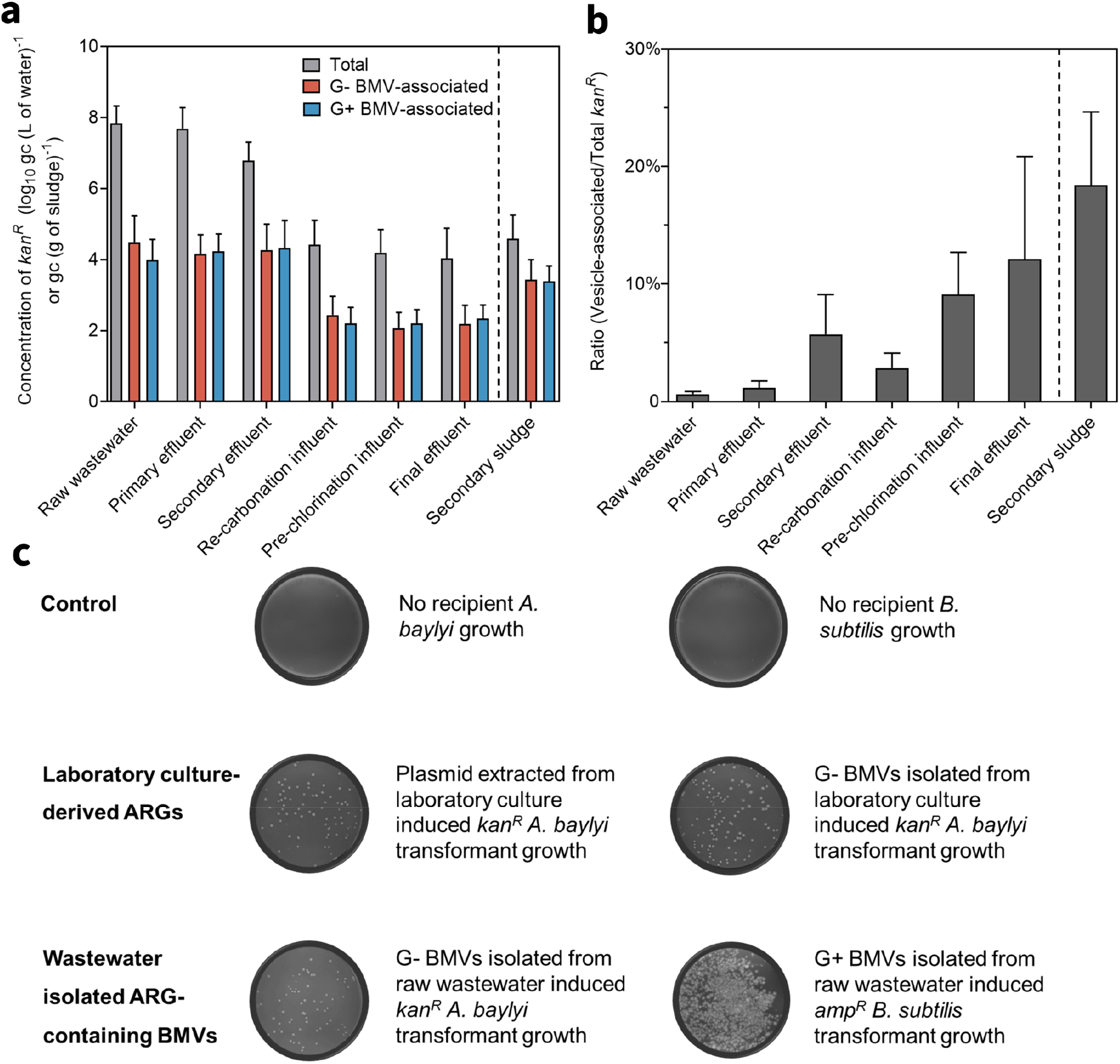
Surveillance of antibiotic resistance genes (ARGs) across the wastewater treatment train and transformation mediated by bacterial membrane vesicle (BMV)-associated ARGs. **a**, Surveillance of total, Gram-negative BMV-associated, and Gram-positive BMV-associated kanamycin resistant genes (*kan*^*R*^) across the treatment train. Gram-negative and Gram-positive are denoted as G- and G+, respectively. Gc represents gene copies. Data are presented as mean ± SD (n=4-8). **b**, Contribution of BMV-associated *kan*^*R*^ to the total pool across the treatment process. Data are presented as mean ± SEM (n=3). **c**, Transformation mediated by BMV-associated ARGs isolated from laboratory cultures and wastewater. Selective agar plates were used to detect antibiotic-resistant bacteria, supplemented with 25 μg mL^−1^ kanamycin or 50 μg mL^−1^ ampicillin.

To assess the role of vesicle encapsulation in ARG persistence, the fraction of *kan*^*R*^ associated with combined Gram-positive and Gram-negative BMVs relative to total *kan*^*R*^ was quantified (**Fig. 2b**). In raw wastewater and primary effluent, it accounted for only 0.63% and 1.18%, indicating most ARGs were intracellular or associated with eDNA/bacteriophages. After secondary treatment, the fraction increased to 5.71%, likely due to degradation of labile ARGs and/or release of BMV-containing ARGs during activated sludge processes. It then decreased to 2.85% after chemical flocculation, suggesting elevated pH and flocculation preferentially removed BMVs. The fraction peaked in pre-chlorination and final effluent at 9.10% and 12.1%, respectively, indicating increasing dominance of vesicle-protected ARGs despite overall reductions in total *kan*^*R*^. BMV-associated *kan*^*R*^ was also enriched in secondary sludge (18.4% versus 5.71% in secondary effluent), likely due to aggregation and hydrophobic interactions promoting accumulation in solids. These results highlight BMVs as a persistent ARG reservoir and underscore the need to evaluate sludge treatment (e.g., anaerobic digestion) for mitigating risks in biosolids reuse.

To determine whether BMV-associated ARGs are functional in promoting HGT, we performed BMV-mediated transformation assays (**Fig. 2c**). Native *Bacillus subtilis* and *Acinetobacter baylyi* ADP1 showed no growth on ampicillin and kanamycin plates, respectively. In contrast, Gram-positive and Gram-negative BMVs purified from raw wastewater by using immunomagnetic isolation (with DNase/RNase treatment) successfully transferred resistance, enabling transformants of *B. subtilis* and *A. baylyi* ADP1 to grow on ampicillin and kanamycin plates, respectively. Wastewater Gram-positive BMVs did not induce growth of *B. subtilis kan*^*R*^ transformants, probably due to host-specific regulatory incompatibility. Positive controls confirmed that both plasmid pIM1522 and *E. coli*-derived BMVs carrying pIM1522 transferred *kan*^*R*^ to *A. baylyi* ADP1, with BMVs showing higher effectiveness (frequency of 8.40 × 10^−5^ versus 1.47 × 10^−5^; efficiency of 1.15 × 10^6^ versus 2.19 × 10^5^ colony forming units μg^−1^). Wastewater-derived BMVs showed lower transformation frequencies (~10^−7^), likely due to damage incurred during wastewater exposure. Overall, these results demonstrate that BMVs are functional vectors for ARG transfer.

### Bacterial membrane vesicles selectively package and reshape ARG profiles during wastewater treatment

Metagenomic analysis revealed that BMV-associated ARG profiles differed from total ARGs. Combining samples from six treatment-stages, total ARGs were dominated by fluoroquinolone, tetracycline, macrolide, peptide antibiotic, and penicillin β-lactam resistance genes, accounting for nearly half of the annotated abundance (**Fig. 3a**). In BMVs, tetracycline, fluoroquinolone, macrolide, and penicillin β-lactam resistance genes remained prevalent, but peptide antibiotic and rifamycin resistance genes were reduced (7.48% and 7.01% in total ARGs versus 3.80% and 3.77% in Gram-negative and 2.40% and 2.48% in Gram-positive BMV-associated AGRs), while monobactam resistance genes increased (3.02% in total ARGs versus 6.35% in Gram-negative and 7.51% in Gram-positive BMV-associated AGRs), with aminocoumarin resistance genes showing similar enrichment. These patterns indicate that ARGs are selectively partitioned into BMVs rather than randomly distributed. This ARG class compositional shift was further supported by alpha- and beta-diversity analyses (**Figs. 3b and 3c**). Compared with total ARGs, BMV-associated ARGs showed lower Shannon diversity and richness in most samples along the treatment train, indicating that BMVs carried a more selective and compositionally constrained subset of the total ARG pool (**Fig. 3b**). Principal coordinates analysis further showed clear separation between total and BMV-associated ARG profiles, and ARG association form significantly explained compositional variation (ADONIS, *p* = 0.001; **Fig. 3c**). In addition, Gram-negative and Gram-positive BMVs also showed distinct ARG profiles, with Gram-negative BMVs enriched in fluoroquinolone, penicillin β-lactam, and cephalosporin resistance genes, and Gram-positive BMVs enriched in tetracycline, macrolide, monobactam, and aminocoumarin resistance genes (**Fig. 3d**). These differences suggest that BMV-associated ARGs are influenced by bacterial origin.

**Fig. 3.**
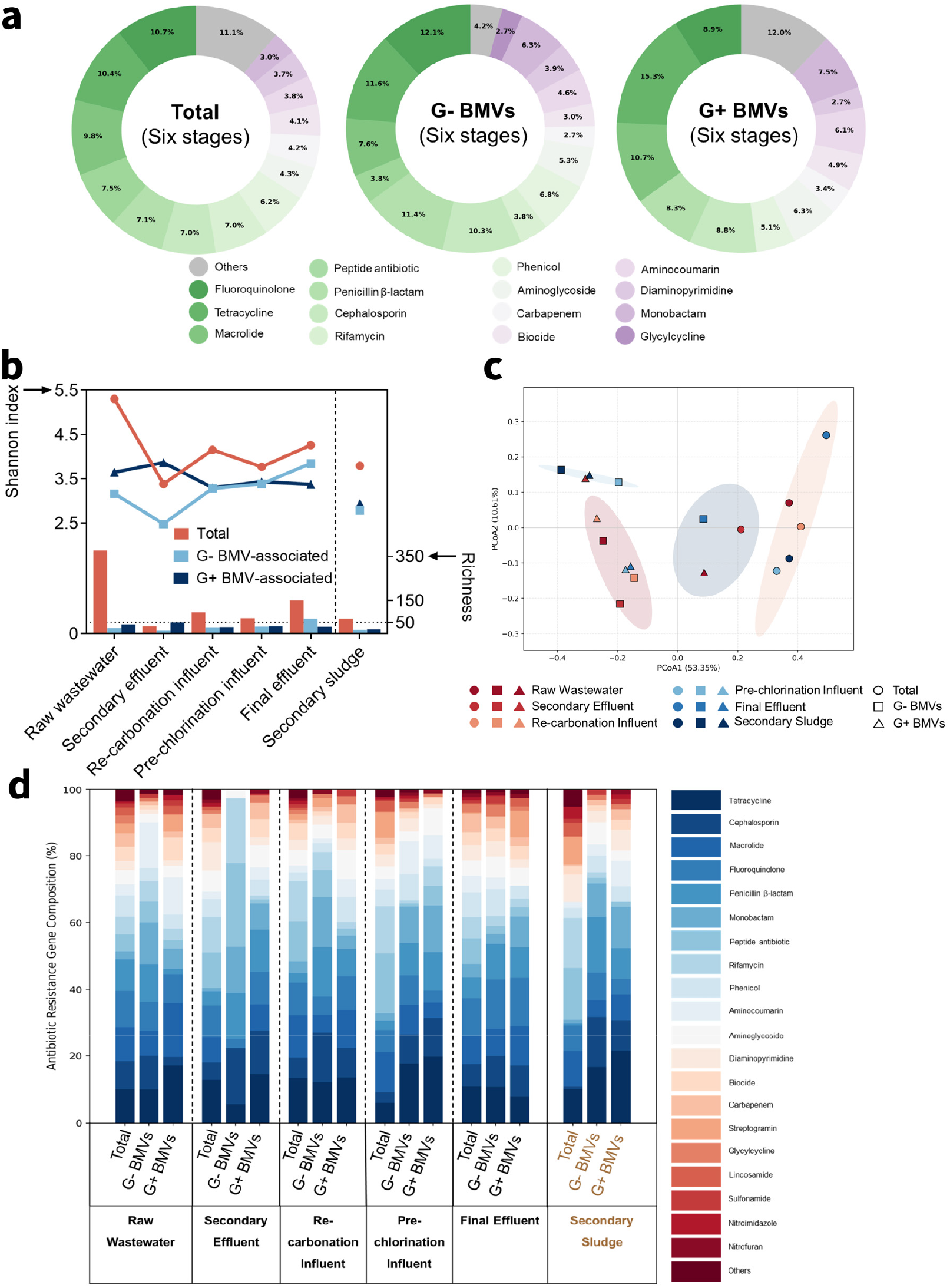
Metagenomic analysis of bacterial membrane vesicle (BMV)-associated and total antibiotic resistance genes (ARGs) across the wastewater treatment train. **a**, Overall ARG class composition pooled across all six treatment-stage samples for total, Gram-negative BMV-associated, and Gram-positive BMV-associated ARGs. ARG classes contributing less than 2.5% were grouped as “Others”. Colors were assigned consistently across samples to represent ARG classes. Gram-negative and Gram-positive are denoted as G- and G+, respectively. **b**, Alpha diversity and richness of ARGs. The lower panel displays ARG richness. The upper panel shows the Shannon diversity index, which reflects both ARG richness and evenness within each sample. A vertical dashed line separates liquid-phase samples from sludge. **c**, Principal coordinate analysis (PCoA) of ARG class composition based on Bray-Curtis dissimilarity. Each dot represents one sample, the color indicates the treatment stage, and the shape indicates ARG-carrying form. The 18 samples were clustered into four distinct groups, each enclosed within a 95% confidence ellipse. **d**, ARG class composition of total, Gram-negative BMV-associated, and Gram-positive BMV-associated ARGs in each treatment step. The top 20 most abundant ARG classes among all the samples are shown, and the remaining low-abundance ARG classes are grouped as “Others”.

Wastewater treatment influenced BMV-associated ARG composition, which did not respond uniformly across treatment steps (**Fig. 3d**). Total ARG profiles shifted during treatment, consistent with previous studies^46,47^; however, BMV-associated ARGs followed a distinct trajectory, indicating a previously underrecognized dynamic. In Gram-positive BMVs, tetracycline, macrolide, and aminocoumarin resistance genes generally decreased from raw wastewater to final effluent, whereas cephalosporin and penicillin β-lactam resistance genes increased, suggesting selective retention or enrichment. In Gram-negative BMVs, aminocoumarin resistance genes declined, while tetracycline, macrolide, cephalosporin, and penicillin β-lactam resistance genes remained relatively persistent; tetracycline and macrolide resistance genes decreased after secondary treatment but reappeared at higher portions in later stages, indicating post-treatment redistribution. Alpha-diversity analysis showed that ARG diversity and richness were highest in raw wastewater and declined sharply after secondary treatment, indicating a substantial narrowing of the detectable ARG pool, with partial recovery in downstream stages suggesting subsequent redistribution of ARGs (**Fig. 3b**). Overall, BMVs represent selective, resilient ARG reservoirs in wastewater rather than passive reflections of the total resistome. Clinically relevant ARGs, such as penicillin β-lactam resistance genes, are preferentially enriched in BMVs and persist through treatment, suggesting that total ARG measurements may underestimate vesicle-mediated dissemination risks.

Our study identifies BMVs as an additional and previously underrecognized carrier of ARGs in municipal wastewater treatment systems, which could highlight several implications for understanding the fate of ARGs in environments. First, by using selective immunomagnetic isolation, we achieved high-purity recovery of Gram-positive and Gram-negative BMVs from complex wastewater matrices and quantified BMV-associated ARGs across treatment. This method provides a framework for understanding the role of environmental and clinical BMVs in bacterial stress adaptation, intercellular communication, and the persistence and dissemination of antimicrobial resistance. Second, the observed higher ratios of BMV-associated to total ARGs and shifts in ARG composition along wastewater treatment indicate selective enrichment and a protective role of BMVs in preserving the encapsulated resistance genes. The BMV-associated ARGs should be considered in microbial resistance surveillance and monitoring, risk assessment, and process design, particularly under increasing water reuse and sustained antibiotic selection pressure. Future work should evaluate how treatment processes such as biodegradation, disinfection, and advanced oxidation affect BMV membrane integrity, cargo stability, and transformation capacity, and whether effective inactivation requires vesicle disruption in addition to reducing total gene abundance.

While this study advances understanding of the dissemination of BMV-associated ARGs in wastewater treatment, several limitations still need to clarify. First, although the immunomagnetic isolation approach enabled selective recovery and quantification of BMVs, further optimization of antibody-biomarker interactions is needed to improve recovery performance in complex matrices and to enhance isolation selectivity. Second, the current workflow is multistep and relatively costly, processing larger volumes or higher amounts of BMVs requires additional antibody input, which may further limit its routine use for large-scale environmental surveillance until simpler and more standardized platforms are developed. Third, although our results support a protective role of BMVs and demonstrate retained transformation activity, the mechanisms by which BMVs protect, stabilize, and deliver ARG cargo remain incompletely resolved. Finally, the impacts of wastewater-derived BMVs beyond the treatment train remain unclear. Investigating their persistence during water reuse, dispersal across environmental media, and interactions with microbial communities, epithelial barriers, and immune systems will be important for assessing their role in antimicrobial resistance dissemination and their potential health risks in wastewater-impacted environments.

## Method

### Bacterial Culture and BMV Production

*Staphylococcus aureus* TCH1516 (BEI Resources NR-10133) hosting a plasmid pUSA300HOUMR, a methicillin resistant variant (MRSA), and *Escherichia coli* (DH5α strain, Addgene, 46012) hosting a plasmid pCM184 were cultured to produce Gram-positive and Gram-negative BMVs, respectively. They were propagated in Luria-Bertani Broth (LB, Fisher Bioreagents, BP1427-500) and Tryptic Soy Broth (Fisher Scientific, DF0370-17-3), both supplemented with 50 μg mL^−1^ kanamycin (Fisher Scientific, BP906-5), in a shaking incubator (200 rpm) at 37 °C for 24 h. Stationary phase cultures were harvested.

50 mL of stock suspension of MRSA and *E. coli* was centrifuged at 14,000 × g for 10 min at 4 °C and subsequently filtered through 0.45 μm mixed cellulose ester (MCE) membranes (Sigma-Aldrich, HAWP04700) to remove large particles and bacterial cells. The filtrate was then ultracentrifuged at 100,000 × g for 3 h at 10 °C, and the pellet containing vesicles was re-suspended in 500 μL of 1× Tris-buffered saline (TBS) containing 0.05% (v/v) Tween 20 (Thermo Fisher Scientific, 28360).

### Wastewater and Sludge Sample Collection

Wastewater and sludge samples were collected from seven treatment stages of a water reclamation facility in Northern Virginia using 24-h composite sampling between May and August 2024. 10 L of conventional treatment samples including raw wastewater influent, primary effluent, and secondary effluent were collected and aliquoted into 1 L each. 2 L of secondary sludge was sampled from secondary clarifiers and aliquoted into 1 L. 100 L of advanced treatment samples, including re-carbonation influent, pre-chlorination influent, and final effluent, was concentrated on-site using a hollow fiber ultrafilter (Dial Medical Supply, ELISIO-25H) operated with a peristaltic pump at 1000 mL min^−1^. The filter was backflushed by 500 mL of 1× PBS supplemented with 3.75 mL of 10% (v/v) Tween 20 to collect all retentate. All samples were stored at 4 °C.

### Pre-treatment of Wastewater and Sludge Samples

1 L of raw wastewater influent, primary effluent, and secondary effluent was firstly filtered with a 20 μm filter paper (Fisher Scientific, 09-795B). Then, 10 mL of 10% (v/v) Tween 20 was added into filtrate. All filtrate was centrifuged at 14,000 × g for 10 min at 4 °C to remove remaining large particles. The supernatant was subsequently subjected to vacuum filtration through a 0.45 μm MCE membrane to remove bacterial cells. Finally, the filtrate was concentrated to a 1.2 mL eluted solution through a concentrating pipette (InnovaPrep). In short, the 0.45 μm membrane filtrate passed through an Ultrafilter Single use Concentrating Pipette Tip (InnovaPrep, CC08003-10), and BMVs accumulated in the retentate were eluted using PBS buffer containing 0.075% (v/v) Tween 20 (InnovaPrep, HC08000) for two times. 1 L of secondary sludge was centrifuged at 20,000 × g for 10 min at 4 °C. After weighing the wet mass, 10 g of collected pellet was re-suspended in 90 mL of PBS and vortexed at 3,000 rpm at room temperature to obtain a sludge suspension. The sludge suspension was filtered through a 0.45 μm MCE membrane, and the filtrate was concentrated to 1.2 mL of eluent using a concentrating pipette. Ultrafiltration retentate of re-carbonation influent, pre-chlorination influent, and final effluent (~ 500 mL) was filtered through a 0.45 μm MCE membrane, and the filtrate was subsequently concentrated by a concentrating pipette to 1.2 mL.

### Immunomagnetic Isolation of BMVs

100 μL of BMV suspension, derived from laboratory culture, wastewater, or sludge, was diluted to 500 μL of reaction volume before the isolation started. For immunomagnetic isolation of BMVs, 20 µg of LTA antibody (Thermo Fisher Scientific, MA5-47519) and 20 µg of lipid A LPS antibody (Bio-Rad, OBT1844) was first incubated with Gram-positive and Gram-negative BMV suspension at 4 °C in a rotator overnight, respectively, forming BMV antigen-antibody complexes. Next, Pierce™ Protein A/G Magnetic Beads (10 mg mL^−1^) (Thermo Fisher Scientific, 88802) were prepared following the user instruction. Briefly, 100 μL of magnetic beads were washed and activated by binding/wash buffer of TBS containing 0.05% (v/v) Tween 20 (Thermo Fisher Scientific, 28360). All the beads were captured by a magnetic stand and all the supernatant was removed. After 2 h incubation, protein A/G magnetic beads were used to capture LTA-bearing Gram-positive BMVs and LPS-bearing Gram-negative BMVs via Fc-mediated binding to the respective antibodies. Purified Gram-negative and Gram-positive BMVs were then harvested in a low-pH elution buffer (Pierce™ IgG Elution Buffer, Thermo Fisher Scientific, 21028) and immediately neutralized by Tris-HCl (pH 8.0, Fisher Scientific, 15-568-025).

### DNase and RNase Treatment

To eliminate potential overestimation caused by extracellular DNA adsorbed onto the surface of BMVs, all BMV samples were subjected to DNase and RNase treatment prior to DNA extraction. The treatment followed the manufacturers’ instructions for Baseline-ZERO™ DNase (Biosearch Technologies, DB0715K) and RNase I (Thermo Fisher Scientific, EN0601). Briefly, 10 molecular biology unit (MBU) of each enzyme were added to 100 µL of vesicle suspension and incubated at 37 °C for 30 min. Enzymatic activity was then terminated by adding a DNase stop solution (Biosearch Technologies, DB0715K), followed by incubation at 65 °C for 10 min to ensure complete inactivation of DNase and RNase.

### DNA Extraction

BMVs isolated from laboratory cultures, wastewater, and sludge are used for DNA extraction and ARG concentration determination. Total ARGs, including those associated with free eDNA, bacteria, bacteriophages, BMVs, and other particles, were extracted from untreated or minimally processed samples. These included raw wastewater, primary and secondary effluents without further treatment; re-carbonation influent, pre-chlorination influent, and final effluent concentrated by onsite hollow fiber ultrafiltration; and secondary sludge that was centrifuged and resuspended in PBS. Details are included in the section of Wastewater and Sludge Sample Collection and Pre-treatment of Wastewater and Sludge Samples. Samples were lysed using PowerWater Bead Pro Tubes (Qiagen, 14900-50-NF-BT) with 5 min of bead beating at a maximum speed (3,000 rpm). High-purity total DNA was extracted with DNeasy PowerWater Kit (Qiagen, 14900-100-NF) according to the manufacturer’s manual and eluted in 30 μl of 10 mM Tris buffer (included in DNeasy PowerWater Kit) for subsequent qPCR quantification and metagenomic sequencing. The quantity and quality of DNA processed with whole-genome sequencing and library construction were assessed by a Qubit 4.0 Fluorometer (Thermo Fisher Scientific) and a TapeStation (Agilent Technologies), respectively.

### qPCR Assay

qPCR was performed on a QuantStudio 3 Real-Time PCR System (Fisher Scientific) to quantify ARGs. Primers and standard templates were obtained from Integrated DNA Technologies and designed to target the kanamycin resistance gene (*kan*^*R*^), ampicillin resistance gene (*amp*^*R*^), tetracycline resistance gene (*tet*^*R*^), β-lactamase coding gene (*bla*^*Z*^), cadmium-binding protein gene (*cad*^*R*^), and crAssphage. Each 20 µL of reaction mixture contained 10 µL of 2× SYBR Green Universal Master Mix (Fisher Scientific, 43-444-63), 1 µL each of forward and reverse primers (0.5 µM final concentration), 1 µL of DNA template, and 7 µL of DNase-free water.

Thermal cycling conditions consisted of an initial denaturation at 95 °C for 10 min to completely activate AmpliTaq Gold DNA Polymerase, followed by 40 cycles of 95 °C for 5 s, 60 °C for 1 min, and 72 °C for 20 s for *kan*^*R*^, *amp*^*R*^, and *tet*^*R*^; 40 cycles of 95 °C for 5 s, 60 °C for 30 s, and 72 °C for 10 s for *bla*^*Z*^ and *cad*^*R*^; and 40 cycles of 95 °C for 5 s and 60 °C for 1 min for crAssphage. Melting curve analyses were conducted with the following steps: denaturation at 95 °C for 15 s, annealing at 60 °C for 15 s, and a gradual increase from 60 °C to 95 °C at 0.1 °C s^−1^ with continuous fluorescence measurement.

Standard curves were generated using 10-fold serial dilutions of the standard templates over six orders of magnitude, analyzed concurrently with unknown samples for absolute quantification. DNase-free water was used as a no-template control. Serial dilutions of extracted DNA from raw wastewater did not indicate any inhibition for qPCR (**Fig. S7**). The qPCR amplification efficiencies ranged from 90% to 110%, with correlation coefficients (R^2^) greater than 0.99. Sequence information was summarized in **Table S1**.

### Recovery of Spiked BMVs from Buffer and Wastewater

*E. coli* BMVs were first purified by immunomagnetic isolation, then spiked into 1× TBS containing 0.05% (v/v) Tween 20 (Thermo Fisher Scientific, 28360) to a final volume of 500 μL and subjected to a second round of immunomagnetic isolation to determine recovery efficiency. Next, MRSA- and *E. coli*-derived BMVs were spiked into particle-depleted wastewater to evaluate recovery of vesicles by immunomagnetic isolation. Particle-depleted wastewater, used to remove endogenous vesicles, was prepared from raw wastewater via sequential centrifugation at 14,000 × g (15 min, 4 °C), 0.45 μm microfiltration (MCE membrane), and ultracentrifugation at 100,000 × g (3 h, 10 °C). Recovery efficiencies of BMVs were determined as the ratio of isolated BMV-associated *cad*^*R*^ or *kan*^*R*^ to its initial spiked concentration. Specifically, *cad*^*R*^ and *kan*^*R*^ were used to track MRSA- and *E. coli*-derived vesicles, respectively.

### Co-isolation of Spiked Plasmid DNA

Plasmids were extracted from MRSA and *E. coli* cultures using the QIAprep Spin Miniprep Kit (Qiagen, 27104) according to the manufacturer’s protocol. The purified plasmids were then spiked into particle-depleted raw wastewater and subjected to the immunomagnetic BMV isolation protocol. Recovery efficiencies of plasmids were determined as the ratio of isolated *kan*^*R*^ or *cad*^*R*^ to its initial spiked concentration. Specifically, *cad*^*R*^ and *kan*^*R*^ were used to track MRSA- and *E. coli*-derived plasmids, respectively.

### Co-isolation of Endogenous Bacteriophages

Unprocessed raw wastewater was subjected to the immunomagnetic BMV isolation protocol, and recovery efficiencies of crAssphage were determined as the ratio of isolated phage to its endogenous concentration in the wastewater.

### Cross-reactivity of Immunomagnetic Isolation

LTA or LPS antibodies were separately incubated with PBS containing BMVs derived from MRSA (Gram-positive) or *E. coli* (Gram-negative), yielding recovery of Gram-positive BMVs via LTA-antibody binding, Gram-negative BMVs via LPS-antibody binding, and cross-binding of Gram-positive BMVs by LPS antibodies and Gram-negative BMVs by LTA antibodies. Recovery efficiencies of BMVs were determined as the ratio of isolated BMV-associated *cad*^*R*^ or *kan*^*R*^ to its initial spiked concentration.

### Transmission Electron Microscopy

15 μL of BMVs isolated from raw wastewater were placed onto a copper grid for 1 min, and excessive sample solution was removed by a filter paper. The sample remaining on the grid was then fixed by 4 wt % glutaraldehyde (Electron Microscopy Sciences) for 5 min in 0.01 M PBS (pH 7.4). The grid was then washed four times with deionized water, each wash lasting 1 min. Subsequently, BMV samples were stained using 5 μL of 1 wt % uranyl acetate (Electron Microscopy Sciences) for 1 min and air-dried overnight to complete staining. TEM images were generated through a FEI Talos™ F200X Transmission Electron Microscope.

### Nanoparticle Tracking Analysis

The hydrodynamic size distribution and particle concentration of raw wastewater and isolated BMVs were measured using a NanoSight NS300 (Malvern Panalytical) following the manufacturer’s instructions. Prior to NTA, concentrated raw wastewater and isolated BMVs were diluted to a final volume of 1 mL with PBS to achieve concentrations within the optimal measurement range. PBS was used as a blank control and to rinse the tubing between measurements. For all measurements, the flow rate, camera level, and detection threshold were kept constant. No significant differences in BMV size distribution were observed before and after Tween 20 addition up to 0.1% indicating that the selected detergent concentration used for pretreating wastewater and sludge did not disrupt vesicles.

### Atomic Force Microscopy

BMVs isolated from laboratory cultures and raw wastewater were first dialyzed in Spectrum™ Spectra/Por™ Float-A-Lyzer™ G2 Dialysis Devices (300 kDa, Spectrum, G235035) against nuclease-free water to remove impurities. Next, BMVs were immobilized on positively charged PLL-functionalized mica for AFM characterizations in liquid. Topographical characterization was conducted in a tapping mode (amplitude-modulation mode) using a BL-AC40TS cantilever (Oxford Instruments). The acquired AFM images were processed by first-order flattening, transverse profile analysis, and three-dimensional reconstruction, and 30 BMVs from each origin were measured for the lateral size and height. Mechanical characterization was then carried out in a contact mode through nanoindentation using an AC160TS-R3 cantilever (Oxford Instruments). The cantilever was rigorously calibrated by measuring the inverse optical lever sensitivity, thermally tuning the spring constant, performing long-range piezo checks, and verifying tip cleanliness. During nanoindentation, the AFM tip was positioned at the center of each BMV and advanced to compress the BMV while recording force-indentation curves. For each BMV, 40 repeated force-indentation curves were acquired. In total, 20 BMVs were characterized mechanically, and the resulting curves were fitted with the Hertz model to estimate membrane stiffness (MPa).

### Transformation Assay

*Acinetobacter baylyi* ADP1 (ATCC 33305) and *Bacillus subtilis* (ATCC 23857) were selected as naturally competent recipient cells for transformation assays using both Gram-positive and Gram-negative BMVs purified from raw wastewater by immunomagnetic isolation and treated with DNase and RNase. Inoculation with laboratory-derived plasmid pIM1522 (pBAV1k-T5-gfp, carrying *kan*^*R*^, Addgene, 26702), as well as Gram-negative BMVs containing pIM1522, served as positive controls. Negative controls without added plasmids or BMVs were also included. Plasmid pIM1522 was selected because it is a shuttle vector compatible with both *E. coli* and *A. baylyi* ADP1, which were used for producing BMVs and naturally competent recipient cells, respectively. BMVs derived from *E. coli* harboring pIM1522 were purified by immunomagnetic isolation and then treated with DNase/RNase (10 MBU per 100 µL of sample). Plasmid pIM1522 was also extracted from these BMVs using the QIAprep Spin Miniprep Kit (Qiagen, 27104). DNA concentration was determined using a Qubit 4 Fluorometer (Thermo Fisher Scientific).

Naturally competent *A. baylyi* ADP1 cells and *B. subtilis* were revived from frozen stocks in 20 mL of LB medium (Fisher Bioreagents, BP1426-500) and incubated at 30 °C for 12-16 h with shaking (200 rpm). To precondition cells, 5 mL of *A. baylyi* ADP1 and *B. subtilis* cultures were next transferred into a 50 mL tube, centrifuged at 6,000 × g for 5 min, washed twice with 1 mL of PBS, and resuspended in 0.5 mL of SOC medium (Teknova, S1660) and fresh LB medium, respectively. Transformation assays were then proceeded by mixing 100 µL of *A. baylyi* ADP1 or *B. subtilis* suspension with 100 µL of BMVs or plasmid DNA and incubated statically at 25 °C for 3 h. Following incubation, mixtures were diluted in LB medium, shaken at 200 rpm for an additional 12-16 h at 30 °C, and plated onto both nonselective agar plates (LB only) and selective agar plates (LB supplemented with 25 μg mL^−1^ of kanamycin or 50 μg mL^−1^ ampicillin). Plates were incubated at 30 °C for 16 h, and colonies were visualized using an iBright 750 Imaging System (Thermo Fisher Scientific). Transformation frequency was calculated as the ratio of transformants on selective plates to total viable cells on nonselective plates, and transformation efficiency was expressed as colony-forming units (CFUs) of transformants per microgram of plasmid DNA.

### Genomics Sequencing

Metagenomic analysis was performed for all samples except primary effluent, as primary treatment was not expected to substantially alter ARG composition^48^ and did not change the concentrations of the selected ARGs in our samples. The DNA samples were first subjected to the whole genome amplification (WGS) by using REPLI-g Mini Kit (Qiagen, 150023) following the manufacturer’s protocol. The sequencing library was then constructed by NEBNext Ultra™ II DNA kit (NEB, E7645S). Pair-end shotgun metagenomics sequencing (2×150 bp) was performed on the Illumina NovaSeq X platform at GENEWIZ, Inc. An average of ~3 Gb of raw reads per sample was obtained. Raw reads were subjected to FastQC (version 0.11.8)^49^ and Trim Glore (version 0.6.6)^50^ for low-quality bases filter, adaptor removal and base trimming. Afterwards, the clean reads were de novo assembled to contigs using MetaSPAdes (version 3.15.5)^51^. Open reading frames (ORFs) were predicted from the assembled contigs using Prodigal (version 2.6.3)^52^ to identify protein-coding genes for downstream ARG annotation. Then, the resulting amino acid sequences were aligned against the Comprehensive Antibiotic Resistance Database (CARD, version 4.0.1)^53^ using DIAMOND (version 2.1.9)^54^ for ARG annotation. To accommodate the fragmented and complex nature of metagenomics DNA extracted from real wastewater samples, E-value <1 × 10^5^ and 60% identity over a query sequence longer than 25 amino acids were set to explore ARGs. The combination of Prodigal and DIAMOND yielded high-confidence protein sequences and enabled rapid protein-protein alignment between samples and the CARD database. The abundance of ARGs was quantified based on the number of hits from DIAMOND alignment and normalized to counts per million reads (CPM) to control for differences in sequencing depth between samples. Identified ARGs were mapped to their corresponding ARG classes based on their annotation in the CARD ontology. ARGs with the same functional classifications were grouped together (e.g., tetracycline antibiotics resistance). The alpha diversity of each metagenomic sample was assessed using the Shannon index (vegan package in R), which considers both the abundance and evenness of ARGs in each individual community across all samples. The observed richness of ARGs further indicates the number of different ARG subtypes detected in each sample. Ultimately, ARG abundance was aggregated and visualized as the distribution of antibiotic resistance categories throughout the entire treatment train. This classification facilitated the interpretation of the resistance mechanisms and microbial risk profiles within the wastewater treatment plant.

### Statistical Analysis

Principal coordinates analysis (PCoA) based on Bray-Curtis distances and ADONIS analysis were carried out with a vegan package in R (v. 4.4.2) assess differences in ARG class composition among samples. Welch’s unpaired two-tailed t-test was applied to AFM-derived parameters (BMV height, lateral size, and membrane stiffness). Differences in ARG abundance measured by qPCR across treatment stages and ARG forms were analyzed using two-way ANOVA, followed by Tukey’s honestly significant difference HSD test. All statistical tests were considered significant at *p* < 0.05.

## Acknowledgments

The project was supported by the United States Environmental Protection Agency grant R840258 and United States Department of Agriculture-National Institute of Food and Agriculture grants 2022-67019-36266 and 2024-67019-42681. We thank the George Washington University Nanofabrication and Imaging Center for performing TEM imaging and the high-performance computing cluster for performing bioinformatic analysis.

