## Supporting Information for "Bacterial Membrane Vesicles in Wastewater Disseminate Antibiotic Resistance Genes"

### Supporting Figures

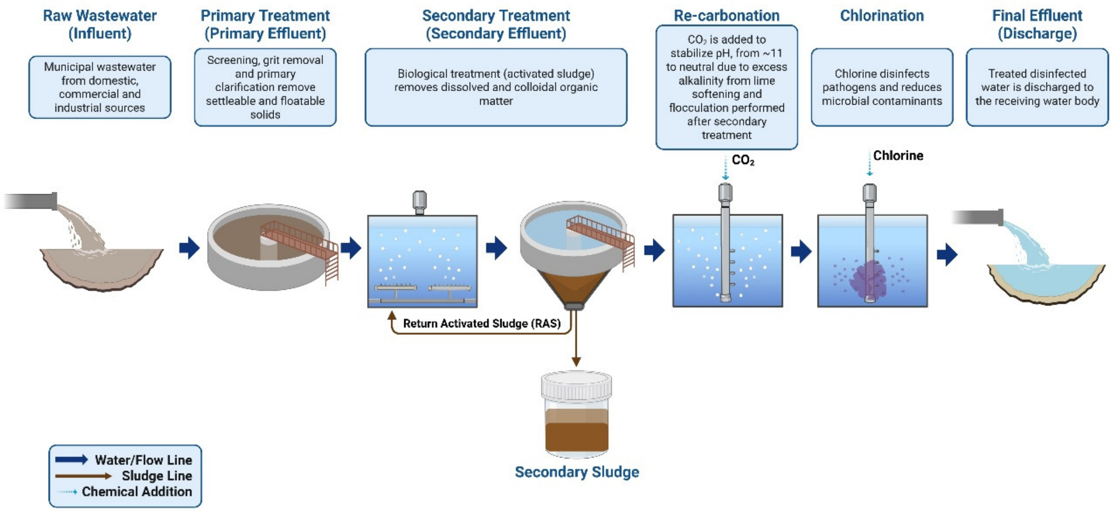


**Figure S1. Schematic illustration of wastewater sampling across the full-scale water reclamation facility.**

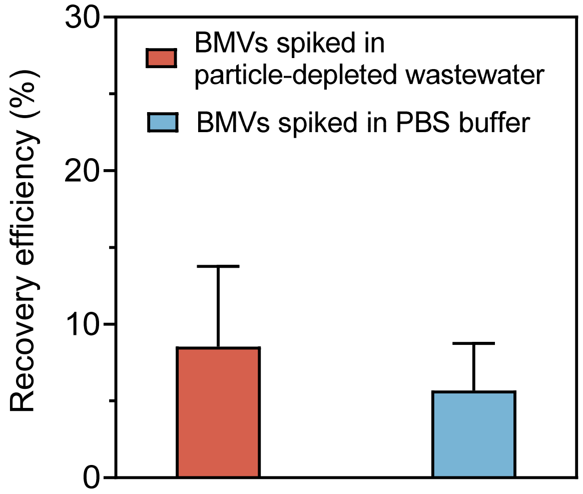


**Figure S2. Recovery efficiencies of BMVs spiked into particle-depleted wastewater and PBS were quantified following ultracentrifugation (100,000 × g, 3 h).** The average recovery efficiencies were 8.53% and 5.67% from particle-depleted wastewater and PBS, respectively. Data are presented as mean ± SD (n=3).

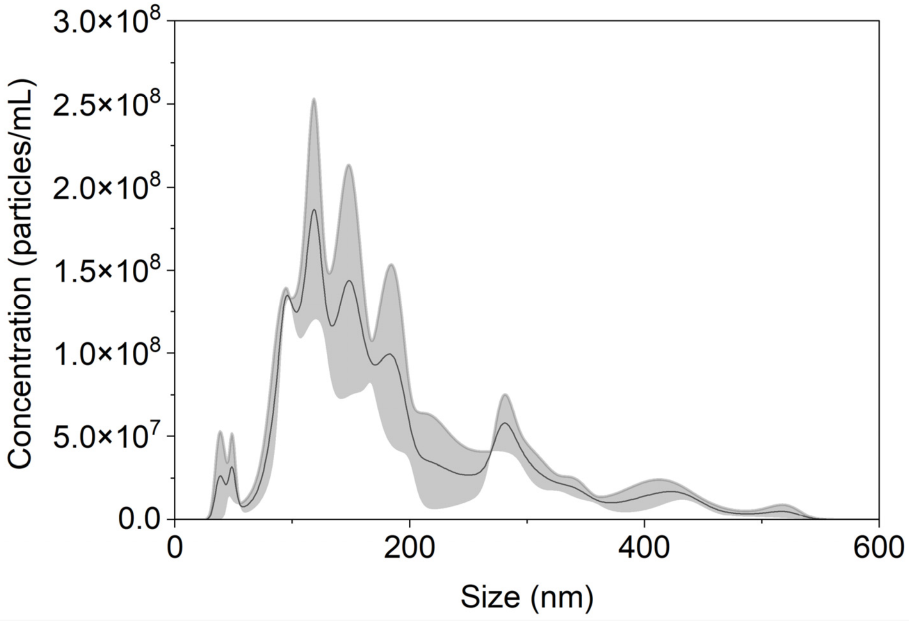


**Figure S3. Characterization of concentrated raw wastewater by NTA.** Concentrated raw wastewater was obtained from raw wastewater after 20 μm filtration, low-speed centrifugation, 0.45 μm filtration, and concentrating pipette concentration (n=3).

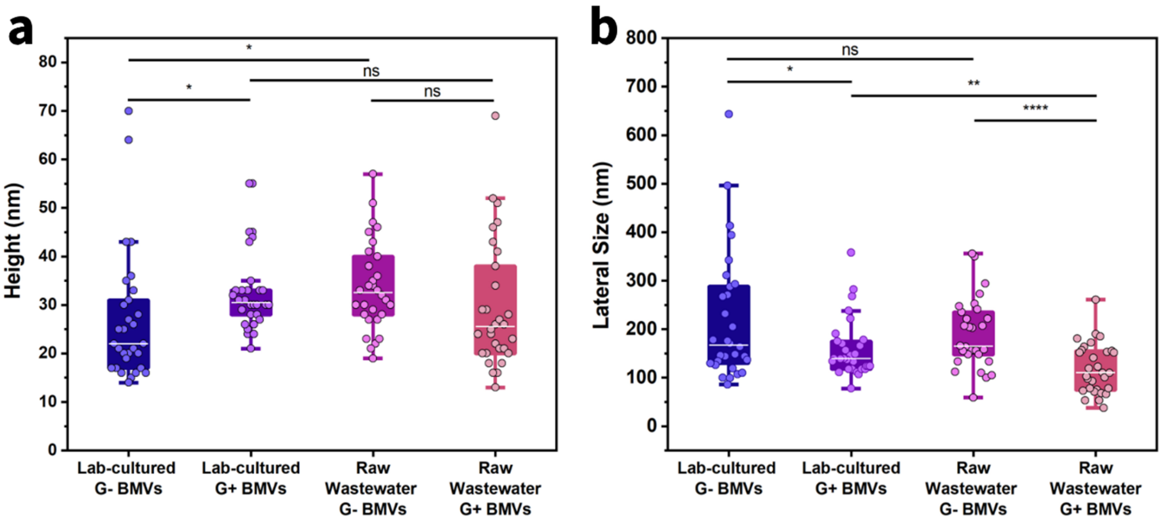


**Figure S4. Dimensions of vesicles isolated from lab-derived bacterial culture and raw wastewater.** (**a**) The height (denoted by *H*) and (**b**) lateral dimension (denoted by *L*) were obtained from the AFM cross-section profile. Each data point represents a *H* or *L* of an individual BMV (n=30). White lines indicate the median, and boxes represent the interquartile range ($25^{th}$-$75^{th}$ percentiles), while whiskers extend to 1.5× the interquartile range from the lower and upper quartiles, with points beyond this range considered outliers if present. Statistical significance is indicated: (ns) $P \geq 0.05$; (*) $0.01 \leq P < 0.05$; (**) $0.001 \leq P < 0.01$; (***) $0.0001 \leq P < 0.001$; (****) $P < 0.0001$.

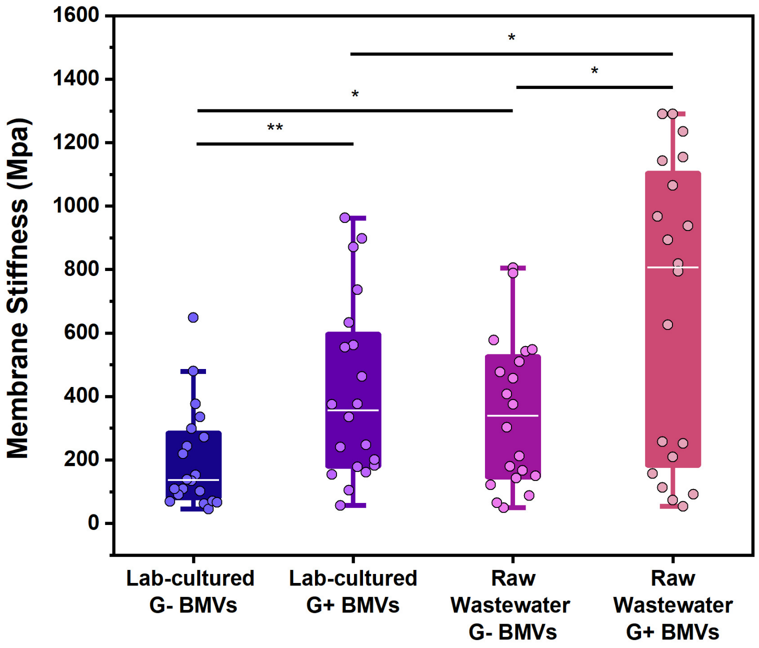

**Figure S5. Membrane stiffness measurement of BMVs by AFM.** Each data point represents the membrane stiffness of an individual BMV (n=20). White lines indicate the median, and boxes represent the interquartile range ($25^{th}$-$75^{th}$ percentiles), while whiskers extend to 1.5× the interquartile range from the lower and upper quartiles, with points beyond this range considered outliers if present. Statistical significance is indicated: (ns) $P \geq 0.05$; (*) $0.01 \leq P < 0.05$; (**) $0.001 \leq P < 0.01$; (***) $0.0001 \leq P < 0.001$; (****) $P < 0.0001$.

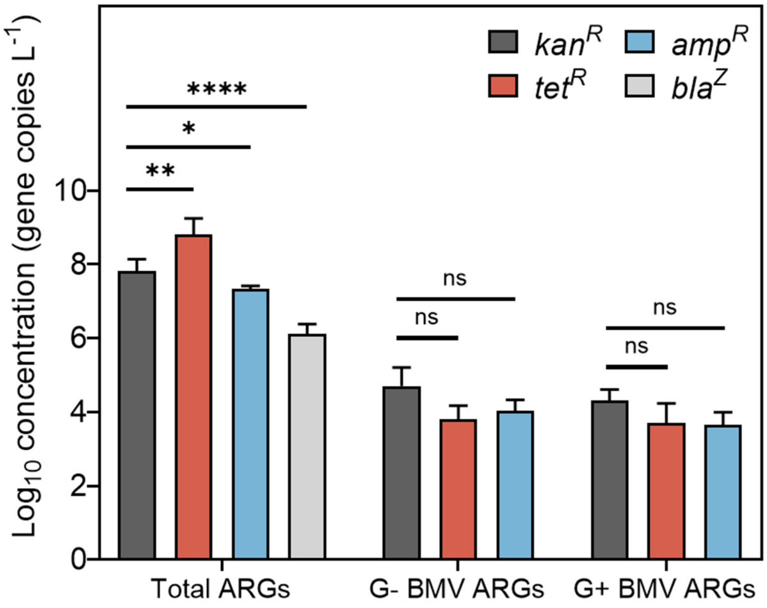


**Figure S6. Quantification of the total and BMV-associated ARG concentrations in wastewater by qPCR.** Error bars represent mean ± SD (n=3-5). Statistical significance between *kan*$^R$ and other ARGs in total and vesicle-associated group is indicated: (ns) $P \geq 0.05$; (*) $0.01 \leq P < 0.05$; (**) $0.001 \leq P < 0.01$; (***) $0.0001 \leq P < 0.001$; (****) $P < 0.0001$.

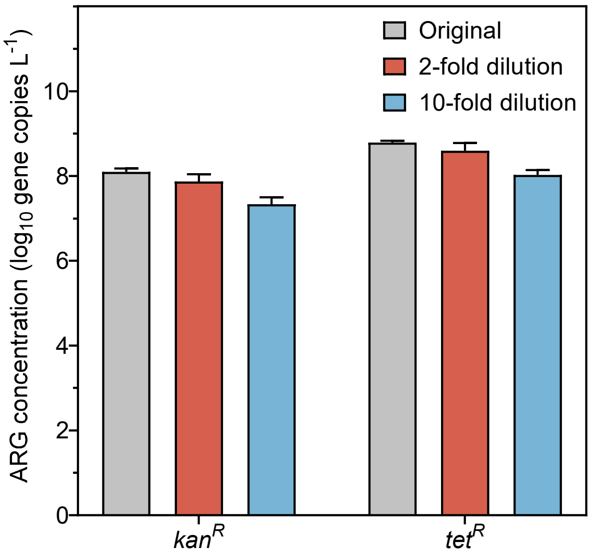


**Figure S7. Serial dilution of DNA extracted from raw wastewater showed no evidence of qPCR inhibition, as assessed using *kan*$^R$ and *tet*$^R$.** Data are presented as mean ± SD (n=3).

**Table S1. Primers and standard templates used for qPCR assay**

| Target | Primer | Standard Template | Reference |
|---|---|---|---|
| Kanamycin resistance gene ($kan^R$) | Forward: TCAACGGGAAACGTCTTGCT<br>Reverse: TCGCGAGCCCATTTATACCC | TAATACAAGGGGTGTTATGAGCCATATTCAACGG GAAACGTCTTGCTCGAGGCCGCGATTAAATTCCAA CATGGATGCTGATTTATATGGGTATAAATGGGCTC GCGATAATGTCGGGCAATCAGGTGCG | 1 |
| Tetracycline resistance gene ($tet^R$)[a] | Forward: GTCACCCTGGATGCTGTAGG<br>Reverse: CCATAGTGACTGGCGATGCT | GTCATCCTCGGCACCGTCACCCTGGATGCTGTAGG CATAGGCTTGGTTATGCCGGTACTGCCGGGCCTCT TGCGGGATATCGTCCATTCCGACAGCATCGCCAGT CACTATGGCGTGCTGCTAGCGCTAT | N/A |
| Ampicillin resistance gene ($amp^R$)[a] | Forward: ACTACGATACGGGAGGGCTT<br>Reverse: TTCCGGCTGGCTGGTTTATT | CGTGTAGATAACTACGATACGGGAGGGCTTACCA TCTGGCCCCAGTGCTGCAATGATACCGCGAGACCC ACGCTCACCGGCTCCAGATTTATCAGCAATAAACC AGCCAGCCGGAAGGGCCGAGCGC | N/A |
| β-lactamase gene ($bla^Z$)[a] | Forward: ACACTCTTGGCGGTTTCACT<br>Reverse: TCCTAAGGGCCAATCTGAACC | ATTCCTTCATTACACTCTTGGCGGTTTCACTTATCA ACTTATCATTTGGCTTATCACTTTTATTGTCTTTAT TCGTAAAAATGACTAAAACAATAGGTTCAGATTG GCCCTTAGGATAAACAAAAGCAAC | N/A |
| Cadmium resistance gene ($cad^R$) | Forward: TGATTGTGAAGGAGAAAAGAGAGC<br>Reverse: GCGCCACAACTTGCTATCG | GGCTATTTATGATGATTGTGAAGGAGAAAAGAGA GCTAAAAAAGAATTGAATGAAAAAGGATTGTCTA AATTAGTTGGTACGGTTGCAATTGTTACGATAGCA AGTTGTGGCGCCGATAATATTGGTTTA | 1 |
| CrAssphage | Forward: CAGAAGTACAAACTCCTAAAAAACGTA GAG<br>Reverse: GATGACCAATAAACAAGCCATTAGC | CAGAAGTACAAACTCCTAAAAAACGTAGAGGTAG AGGTATTAATAACGATTTACGTGATGTAACTCGTA AAAAGTTTGATGAACATACTGATTGTAATAAAGCT AATGGCTTGTTTATTGGTCATC | 2 |

Note: N/A represents not applicable. a: The primers were designed using the Primer-BLAST tool on the NCBI platform.
